# Transcriptional Analysis of *Colletotrichum fructicola* from Different Geographic Regions Inoculated to *Camellia oleifera*

**DOI:** 10.1101/2020.07.13.198457

**Authors:** Shimeng Tan, Yanying Chen, Guoying Zhou, Junang Liu

## Abstract

Anthracnose, caused by Colletotrichum. Spp., is an important disease affecting oil tea (*Camellia oleifera*). Extensive molecular studies have demonstrated that *Colletotrichum fructicola* is the dominant pathogen of oil tea anthracnose in China. The study aims to investigate differences in molecular processes and regulatory genes at different stages of infection of *C. fructicola*, to aide in understanding differences in pathogenic mechanisms of *C. fructicola* of different geographic populations. Materials and Methods: We compared the pathogenicity of *C. fructicola* from different populations (Wuzhishan, Hainan province and Shaoyang, Hunan province), and gene expression of representative strains of the two populations before and after inoculation in oil tea using RNA sequencing. In results we found that *C. fructicola* from Wuzhishan has a stronger ability to infect and impact oil tea leaf tissue. Up-regulated genes in oil tea following infection with the two geographic populations were associated with a number of ribosome-related GO terms and ribosome metabolic related KEGG pathways, and were significantly enriched in galactosidase activity, glutamine family amino acid metabolism, arginine and proline metabolism. In addition, up-regulated gene lists associated with infection in the Wuzhishan strains were significantly enriched purine metabolism pathways as well, while Shaoyang strains were not significantly enriched in these processes. Differences in gene expression were indicative of greater active transcription and translation in the Wuzhishan population while more active glycoprotein metabolism was observed following inoculation with strains from Shaoyang. These results indicate that *C. fructicola* obtained sugars and amino acids from oil tea tissue to resist host immune pressure. Moreover, more active in transcriptional and translational activity and the greater regulation of purine metabolism pathway in the Wuzhishan strain might contribute to its stronger pathogenicity.

## 1 Introduction

Oil tea (*Camellia oleifera*) is an agronomically and culturally important edible woody oil tree species found in China. Oil extracted from the seeds of *oil tea* is comparable to olive (*Olea europaea*) oil. *Oil tea* is widely distributed between the northern latitudes of 32°57’ and 11°17’ in China (Cui, X. et al, 2016). Suitable climates for growth of *oil tea* are generally mild, receive sufficient sunshine and rain, and have short or no ice periods, therefore occurrence of disease and damage due to insect pests are frequent (Zhou, W. et al, 2017). Anthracnose is a prevalent disease which affects *oil tea*, is caused by *Colletotrichum fructicola*, and is readily dispersed via spores. Spores of anthracnose can survive winter conditions and result in continuous infections year over year (Ye Jian-ren, H. W., 2011). Anthracnose can cause leaf fall, flower fall and fruit damage, which can impact yield or oil quality, leading to economic loss.

Colletotrichum fungus is a globally distributed and important pathogen. Colletotrichum fungus is a hemibiotrophic plant fungus, and can affect almost all crops and economic plants. *Colletotrichum* spp. is often spread as conidia though wind, rain, insects. Conidia germinate in aqueous environments to form appressoriums and produce infection nails which can puncture epidermal tissues such as plant leaves and peels. After invading the host, mycelia of colletotrichum fungi are quickly formed between plant cells, destroy plant tissues and use their nutrients for further replication (Chen, X., et al, 2009, Wang, H. K., et al, 2004). To date, 8 species of anthracnose pathogens of *oil tea* have been classified based on morphological and polygenic molecular identification (Jiang, S. Q., et al, 2018, Li, H., et al, 2017, Tang, Y. L., et al, 2015). *C. fructicola* has the highest isolation rate in diseased oil tea leaf samples in China, and is the dominant pathogen of *Camellia oleifera* anthracnose (Li, H., 2018). Recent research on the prevention and treatment of anthracnose in oil tea has focused on application of new chemicals or biological controls such as bacteria (Deng, L., et al, 2017, Zhou, Y. H., et al, 2015). However, it has been suggested that to prevent occurrence of anthracnose in oil tea, it is necessary to have a deeper understanding of its pathogenic mechanisms to develop specific control measures. Therefore, in the current study we focused on the pathogenesis of *C. fructicola* on oil tea leaves, with the aim of improving our understanding of the molecular mechanisms of pathogenesis of *C. fructicola* infection of oil tea.

When a pathogen colonizes and infects a plant, defense responses aim to nutritional starve the fungi and subject it to reactive oxygen species (ROS) stress (Deng, Y. Z., Naqvi, N. I., 2019). Initiation and control of plant defense responses rely on flexible and rapid molecular regulation of numerous pathways in responses to incursion by a pathogen. The infection of plants by hemibiotrophic vegetative fungi is a continuous and staged process whereby infection of plants by *Colletotrichum* fungi can be divided into two stages: vegetative and death phase. Hyphae growth of fungi and its spread is dependent on nutritional availability (Kenneth J, C., 2002). Transcriptome sequencing has emerged as a useful tool for investigating the molecular regulation of different stages of infection by *C. fructicola*.

To investigate molecular mechanisms of pathogenicity of *C. fructicola* from different geographical populations, we analyzed fungi isolated from oil tea samples produced in Shaoyang, Hunan province and Wuzhishan, Hainan province, China via RNA sequencing.

## 2 Materials and Methods

*C. fructicola* used for experiments was isolated from diseased oil tea leaves collected from Shaoyang, Hunan and Wuzhishan, Hainan, which were stored in freezing tube containing 30% glycerin aqueous solution at −80 °C. Four strains of *C. fructicola* were from Shaoyang, Hunan, and seven strains of *C. fructicola* for test were from Wuzhishan, Hainan. The geographical and climatic characteristics of the two locations are shown in Table 1 (Ma, W., 2010, Su, F., 2018, Zheng, D. J., et al, 2016).

**Table 1.**
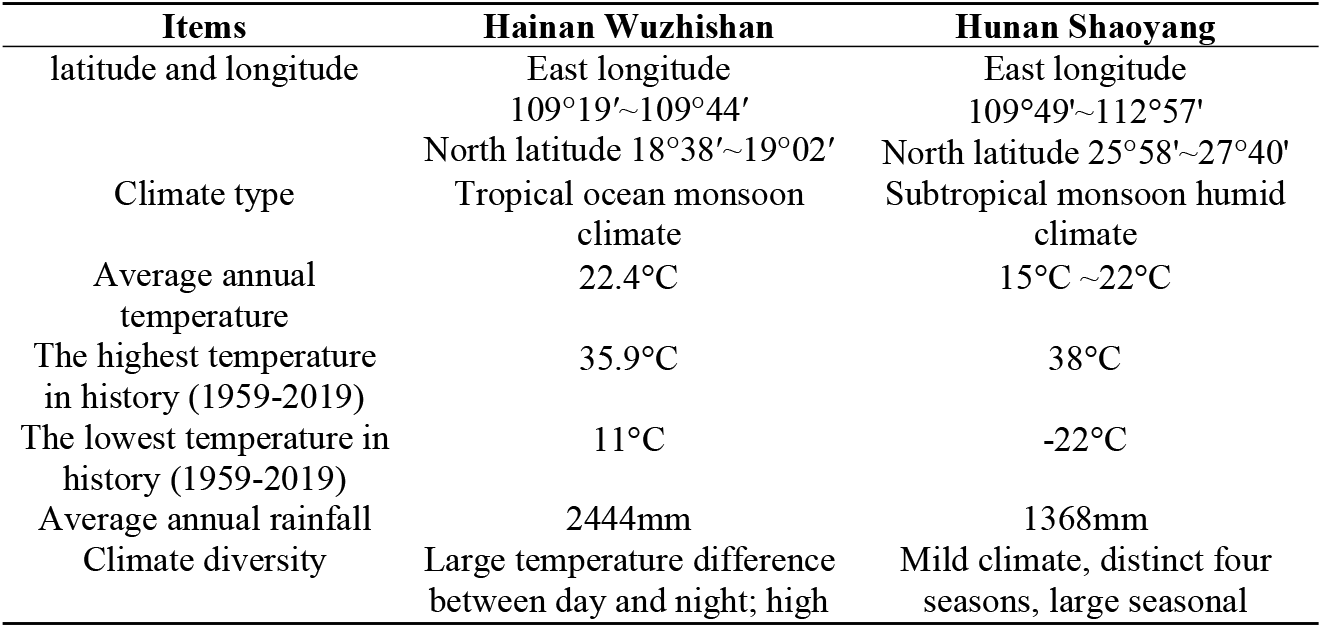
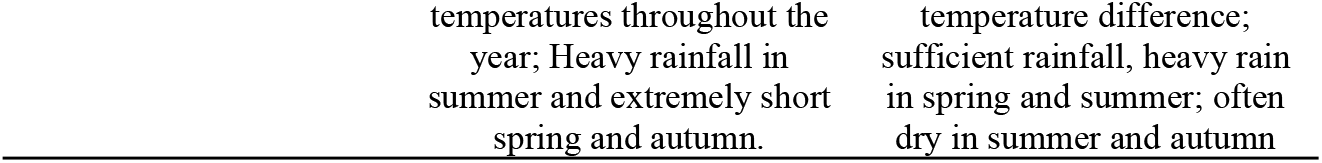
Geographical and climatic features of Wuzhishan and Shaoyang.

| Items | Hainan Wuzhishan | Hunan Shaoyang |
| --- | --- | --- |
| latitude and longitude | East longitude<br>109°19'~109°44' | East longitude<br>109°49'~112°57' |
|  | North latitude 18°38'~19°02' | North latitude 25°58'~27°40' |
| Climate type | Tropical ocean monsoon<br>climate | Subtropical monsoon humid<br>climate |
| Average annual<br>temperature | 22.4°C | 15°C ~22°C |
| The highest temperature<br>in history (1959-2019) | 35.9°C | 38°C |
| The lowest temperature in<br>history (1959-2019) | 11°C | -22°C |
| Average annual rainfall | 2444mm | 1368mm |
| Climate diversity | Large temperature difference<br>between day and night; high | Mild climate, distinct four<br>seasons, large seasonal |

### 2.1 Media Preparation

Potato dextrose agar medium (PDA medium) consisted of; 200g peeled potatoes (from local market in Changsha city, China) was added to 1000mL water and heated for 20 minutes. The mixture was filtered through gauze, and 20g glucose and 20g agar were added to the filtrate. The mixture was autoclaved at 121 °C for 20 minutes. The potato dextrose broth medium (PDB medium) was pruduced without agar. One gram of yeast extract was added to 1L of PDA or PDB medium to promote production of conidia.

### 2.2 Inoculation Experiment on Leaves of Oil Tea

The pathogenicity of the tested strains was determined *in vitro* by the injury inoculation test using leaves of the *Oil tea*, Huashuo. Young leaves were collected, petioles were sealed with paraffin wax, and then placed into a glass petri dish with wet cotton pads added to maintain humidity. A sterilized needle was used to make six pinholes along each leaf surface, three on left half and three on right half of each leaf. There would be eight leaves replicates and 48 pinholes in total. Leaves were inoculated with mycelium from Shaoyang and Wuzhishan strains, the left half with Shaoyang and right half of the leaf with Wuzhishan, each pinhole was inoculated with a 6mm diameter mycelium piece, for 24 inoculation samples per strain total.

Petri dishes were sealed with parafilm and cultured in the dark at 28 °C for 96 hours. Leaves were then removed and the diameter of the diseased spots was measured and photographed.

### 2.3 Sampling of RNA

Two representative strains from Wuzhishan (WZS) and Shaoyang (SY) were cultured on a PDA solid medium for 96 hours. Hyphae pieces at the edge of several colonies were cut and transferred to 100 mL of prepared PDB medium, and cultured on a shaker at 160 rpm and 28 °C for 48 hours. The culture solution was filtered through three layers of microscope wipe paper into new centrifuge tubes and then centrifuged at 5000 rpm for 3 minutes. The supernatant was poured off, spores were rinsed twice with sterile water and sterile water was added to reconstitute spores at a concentration of 1 × 10^9^ pcs/mL. Next, 1.5 mL of spore solution was placed into a 1.5 mL centrifuge tube and centrifuged at 12000rpm for 2 min. The supernatant was removed to obtain conidia samples which were immediately frozen with liquid nitrogen and stored in a −80 °C refrigerator.

Sterile water was added to the remaining spore fluid for dilution, and the number of spores was counted to adjust the concentration of spore fluid to 1 × 10^6^ pcs/mL. Petioles were sealed with wax and placed in a petri dish with wet absorbent cotton. The spore fluid was evenly sprayed onto the leaves, and then the petri dish was sealed. Leaves were incubated at 28 °C in the dark for 96 hours. After incubation, leaves were cut and placed into collection tubes, then immediately frozen with liquid nitrogen and transferred to a −80 °C refrigerator. There were 3 biological replicates per sample.

### 2.4 RNA Sequencing and Data Analysis

The spores and lesion samples were sent to Genedenovo Biotechnology Co., Ltd, Guangzhou, China. for RNA extraction and RNA sequencing. Among them, spore sample group number is identified as S (Start, the point of 0 hpi (hour post inculation), and lesion sample group number is L (Later, the point of 96 hpi). Data analysis was performed online using the Omicsmart platform (Genedenovo Biotechnology Co., Ltd, Guangzhou, China; https://www.omicsmart.com/).

### 2.5 Real-time PCR

The kits: FastQuant RT Kit with gDNase (TIANGEN Biotech Co., Ltd, Beijing, China); SuperReal PreMix Plus (SYBR Green) (TIANGEN Biotech Co., Ltd, Beijing, China) were used for analysis of samples.

Samples of RNA subjected to RNA-seq analysis used the FastQuant RT Kit with gDNase (TIANGEN) for first-strand cDNA synthesis. Five sequences for RNA-seq consistency test were randomly selected from the differentially expressed gene data obtained by RNA sequencing, and twelve sequences for unique up-expression consistency test were from the gene data obtained by RNA sequencing of Wuzhishan strains. All the quantitative PCR primers were designed using NCBI primer-blast. Primer information is shown in Table 2. Primer synthesis was completed by Beijing Qingke Biotechnology Co., Ltd. The Actin gene, which was stably expressed in all RNA-seq samples and was highly expressed, was selected as the reference gene (He, L., et al, 2016). The real-time PCR test uses SuperReal PreMix Plus (SYBR Green). The reaction system consisted of 20 L, including 1 g of cDNA template (100 ng/L), and 0.75 L for each of the front and back primers. Data analysis was performed using QuantStudio ™ Design & Analysis Software (version 1.5.1, Thermo Fisher Scientific). For qRT-PCR data, relative expression log_2_FC was calculated using the Ct method, and compared with RNA sequencing data. Each sample had three replicates.

**Table 2.**
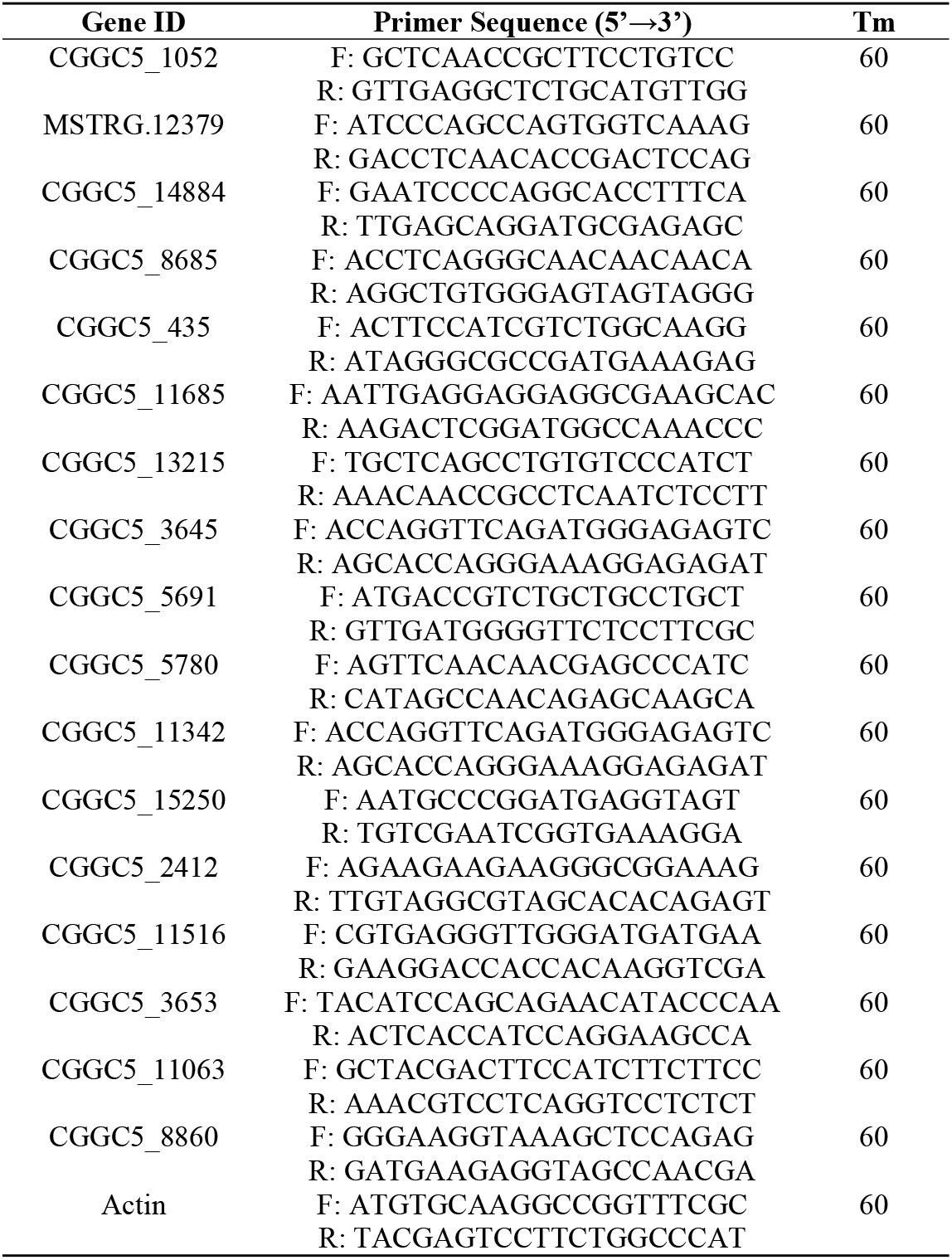
Primer related information for real-time PCR.

### 2.6 Statistical Analysis

The diameters of *C. fructicola* strains from Shaoyang population and Wuzhishan population were counted after 96 hours of cultivation, and the average diameter of the *C. fructicola* from two populations was calculated using IBM SPSS Statistics 20 (IBM, U.S.A). The diameter of *C. fructicola* from two populations were compared by independent sample t-test and two-sided test (α = 0.05).

## 3 Results

### 3.1 Comparison of Mycelia Pathogenicity of Two Populations

Diameter of lesions following inoculation was counted and results are shown in Figure 1. Average diameter of lesions following incubation with the Shaoyang population was 0.37 cm, and for the Wuzhishan population was 0.56 cm. An independent sample T test was performed on lesion diameter among the two populations, and differences were significant (P=0.046 <0.05).

**Figure 1.**
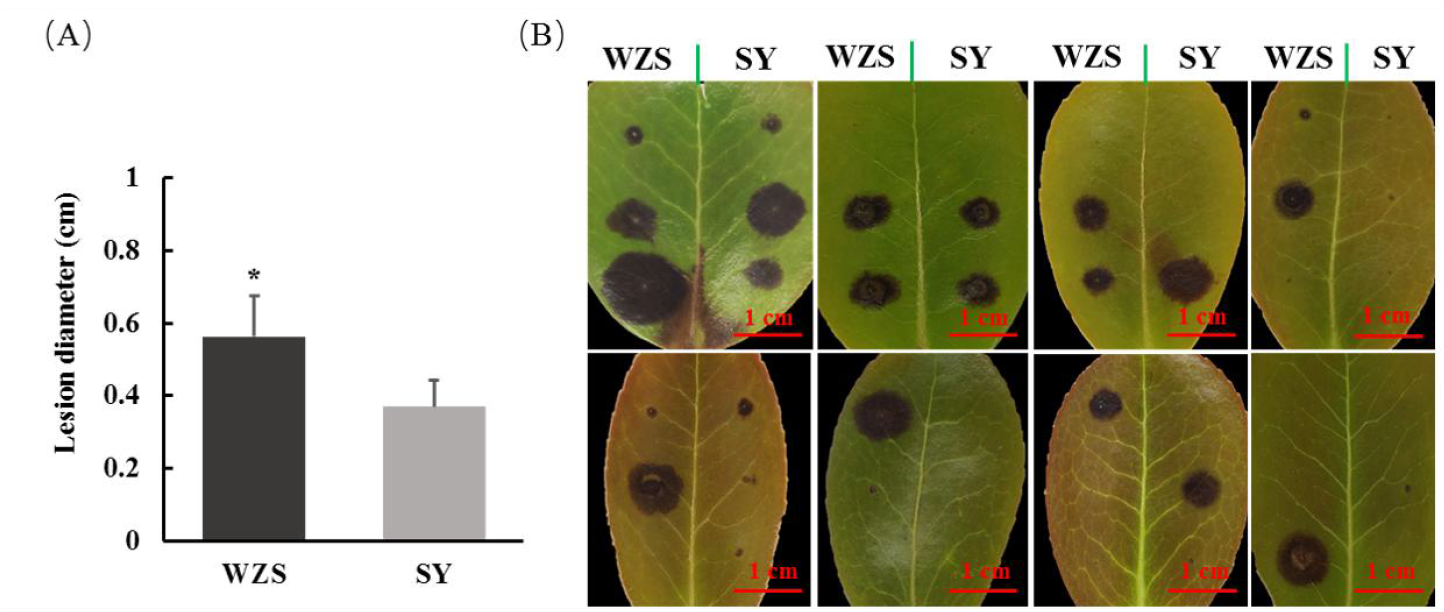
Pathogenicity of *C. fructicola* populations collected from Shaoyang and Wuzhishan to oil-tea leaves. A: Lesion diameters 96 hpi on oil-tea leaves following inoculation with *C. fructicola* populations from Wuzhishan and Shaoyang. Error bars are mean ± standard deviation and asterisks represent significance at *P<0.05(P=0.046). B: Lesions 96 hpi. Green vertical lines represent leaf veins. Wuzhishan strains and Shaoyang strains were inoculated on the left and right sides of leaves.

### 3.2 Comparison of Pathogenicity within Populations and Selection of Representative Strains

In order to select a representative strain from each of the two populations for transcriptome analysis, a comparative test of pathogenicity among the strains in the two populations was performed.

Results of the pathogenicity test of Wuzhishan populations are shown in Figure 2A. After 96 hours of inoculation, average diameter of lesions caused by 7 strains of Wuzhishan-origin was 0.50 cm (The dotted line in Figure 2A). Among the seven strains tested, WZS0202b was the weakest pathogen, whereas WZS0402a and WZS0203b had the highest pathogenicity.

**Figure 2.**
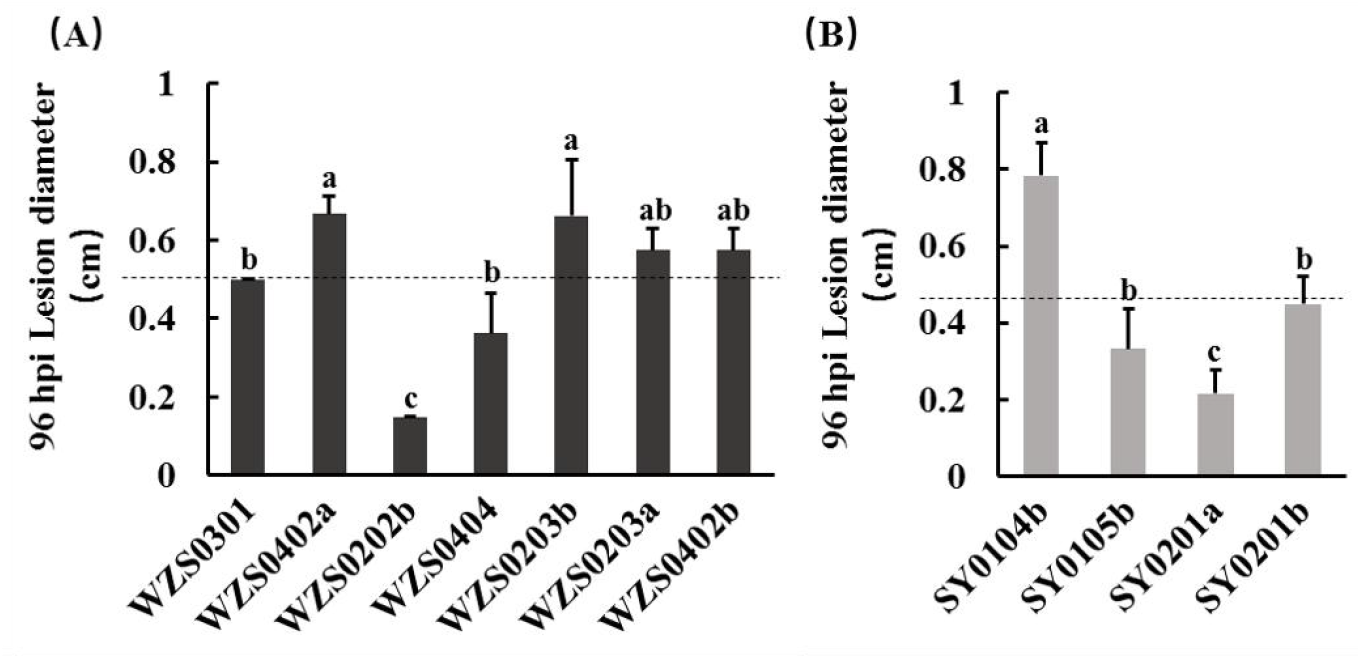
Pathogenicity of C. fructicola strains. A: Pathogenicity of 7 C. fructicola strains from Wuzhishan. The dotted line is the average lesion diameter. Error bars are mean ± standard deviation. B: Pathogenicity of 4 C. fructicola strains from Shaoyang. The dotted line is the average lesion diameter. Error bars are mean ± standard deviation.

Results of the pathogenicity test of Shaoyang populations are shown in Figure 2B. After 96 hours of inoculation average diameter of lesions was 0.45cm (The dotted line in Figure 2B). The pathogenicity of the strain SY0104b was the highest, while pathogenicity of SY0201a was the weakest.

Based on results of the two strains of *C. fructicola*, WZS0402a and SY0104b were selected for RNA sequencing.

### 3.3 Real-time PCR

To verify reliability of sequencing data, five differentially expressed genes were randomly selected for real-time PCR. The gene Actin was selected as the reference gene. Quantitative PCR results showed that except the gene CGGC5_435 in SY-S-vs-SY-L, most gene expression trends were consistent with transcriptome data (Figure 3). In addition, relative expression levels (log_2_FC value) were not significantly different from those of the RNA sequencing results, indicating that transcriptomic sequencing results are reliable.

**Figure 3.**
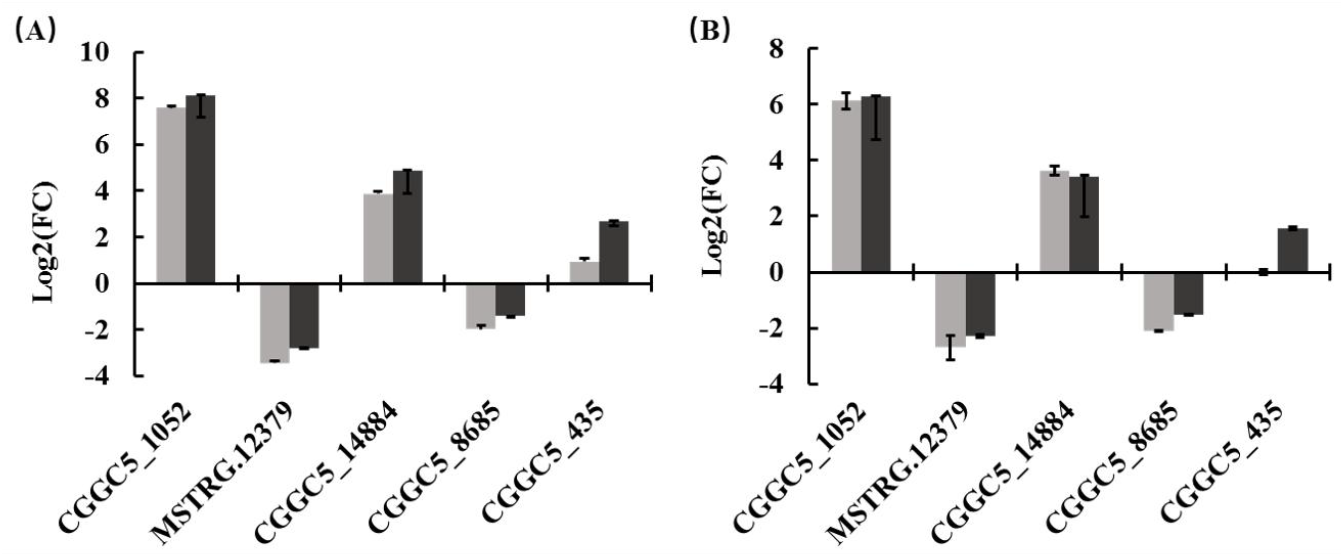
Gene expression of qRT-PCR and RNAseq. A: qRT-PCR and RNAseq verification of WZS-S-vs-WZS-L. B: qRT-PCR and RNAseq verification result of SY-S-vs-SY-L.

To verify reliability of Wuzhishan strains unique expression, twelve differentially expressed genes were selected for real-time PCR. The gene Actin was selected as the reference gene. Quantitative PCR results showed that most selected gene expression trends were consistent with transcriptome data (Figure 6). Furthermore, relative expression levels (log_2_FC value) were not significantly different from those of the RNA sequencing results, revealing that the unique up-expression results of Wuzhishan strains are reliable.

### 3.4 Statistics of Differentially Expressed Genes

Differentially expressed genes were screened using FDR (False Discovery Rate) ≤ 0.05 and |log_2_FC| ≥ 1 as thresholds, and then genes were manually eliminated if they had low expression levels (FPKM <0.5), resulting in selection of WZS-S-vs-WZS-L, SY-S-vs-SY-L. Using homologous spore samples as a control, the number of genes up- and down-regulated in the infection groups were tabulated (Figure 4A). In the WZS-S-vs-WZS-L group, a total of 7846 differential genes were screened, 5951 of which expression was up-regulated (75.85%), and 1895 (24.15%) down-regulated. In the SY-S-vs-SY-L group screening, a total of 7682 differential genes were identified, 5280 (68.73%) of which were up-regulated, and 2402 (31.27%) were down-regulated. During the interaction stage of oil tea infected with *C. fructicola*, the number of up-regulated genes was significantly higher than that of down-regulated genes in the two representative strains. The number of up-regulated genes of *C. fructicola* from Wuzhishan after infection was higher than that from the Shaoyang strains.Comparison of differentially expressed gene lists post-inoculation demonstrated that 4089 genes were up-regulated in both strains after 96 hours of infection (Figure 4B). Overall, similarly expressed genes accounted for 68.7% (WZS) and 77.4% (SY) of up-regulated genes, respectively.

**Figure 4.**
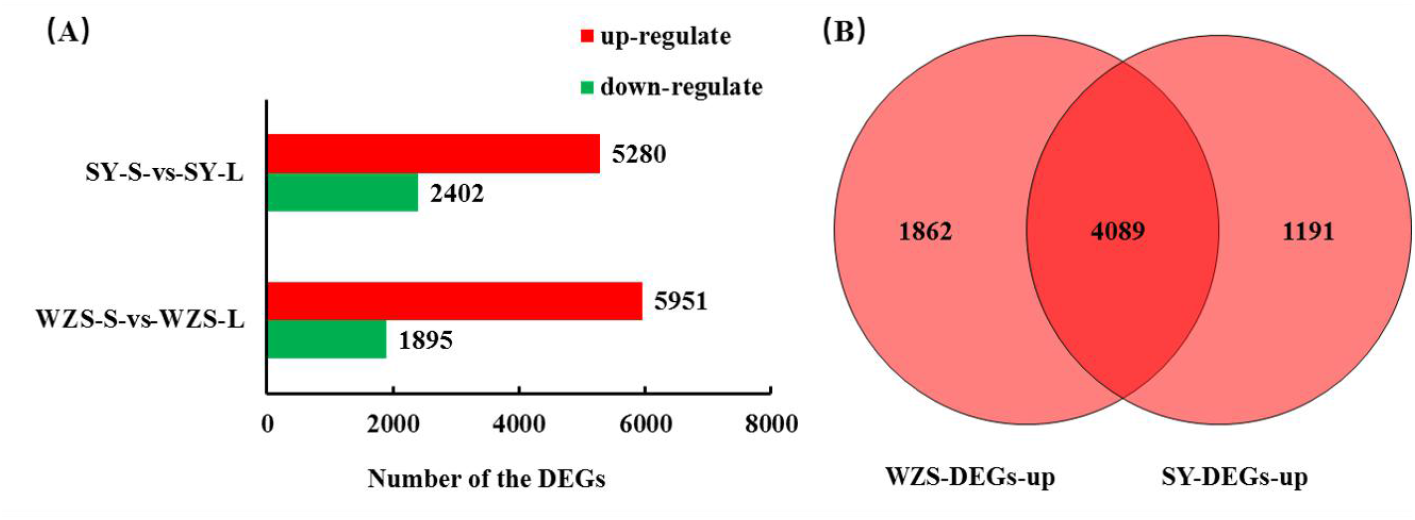
Overview of differentially expressed genes. A: Comparison of differential genes before and after infection. B: Venn diagram of number of up-regulated genes after infection.

### 3.5 Gene Ontology Enrichment Analysis

Gene ontology (GO) function enrichment analysis was performed on differentially expressed genes following infection of WZS-S-vs-WZS-L and SY-S-vs-SY-L groups. As shown in Table 3, GO terms were screened with Pvalue ≤ 0.01 as the threshold. It was found that inoculation with the representative strain of Wuzhishan resulted in enrichment of preribosome, large ribosomal subunit and organelle ribosome GO terms. Identified molecular functions were mainly related to structural molecular activity, translation factor activity (RNA binding), and galactosidase activity. Biological processes were mainly related to the glutamine family amino acid metabolism process.

**Table 3.** GO terms significantly enriched in oil tea leaves inoculated with strains of *C. fructicola* and related up-regulated genes.

| GO ID | Term | All genes with GO annotation | DEGs with GO annotation |  | Pvalue |  |
| --- | --- | --- | --- | --- | --- | --- |
|  |  |  | WZS | SY | WZS | SY |
| GO:0030684 | preribosome | 88 | 84 | 58 | 6.17E-26 | 1.42E-07 |
| GO:0015934 | large ribosomal subunit | 38 | 37 | 28 | 9.39E-13 | 1.08E-05 |
| GO:0000313 | organellar ribosome | 48 | 44 | 36 | 2.61E-12 | 2.71E-07 |
| GO:0005198 | structural molecule activity | 61 | 49 | 37 | 1.36E-10 | 8.63E-05 |
| GO:0008135 | translation factor activity, RNA binding | 65 | 42 | 23 | 5.36E-05 | 0.6050 |
| GO:0015925 | galactosidase activity | 20 | 16 | 15 | 3.25E-04 | 4.76E-04 |
| GO:0009064 | glutamine family amino acid metabolic process | 51 | 32 | 31 | 1.55E-03 | 5.61E-04 |
| GO:0009100 | glycoprotein metabolic process | 30 | 11 | 18 | 0.7554 | 9.73E-03 |

Screening of GO enrichment results of the Shaoyang group, demonstrated that enriched cell composition terms were related to ribosome precursors, large ribosome subunits, and organelle ribosomes. Molecular functions were mainly related to structural molecular activity and galactosidase activity. Identified biological processes were mainly related to amino acid metabolic processes and glycoprotein metabolic processes of the glutamine family.

As presented in Table 3, it can be seen that cell composition terms for the two groups were significantly enriched in ribosomal-related annotations, but the number of differentially expressed genes in the corresponding GO annotations in the Shaoyang group was significantly lesser than that in the Wuzhishan group. Enriched molecular functions among the two groups were related to structural molecular activity and galactosidase activity, however RNA-bound translation factor activity that was enriched in the Wuzhishan group was not significantly enriched in the Shaoyang group. Enriched GO biological processes in both groups were related to glutamine amide family amino acid metabolism. However, glycoprotein metabolism was enriched in the Shaoyang group and not in the Wuzhishan group. Of the two co-enriched annotations, genes associated with galactosidase activity and amino acid metabolism of the glutamine family were most similar.

### 3.6 KEGG Pathway Enrichment Analysis

Analysis of gene lists identified 7 KEGG pathways (Q value ≤0.01) following inoculation with the Wuzhishan and Shaoyang populations (Table 4). Of identified GO pathways, ribosome (ko03010), arginine and proline metabolism (ko00330) were significantly enriched in both groups, indicating that arginine and proline metabolism play an important role in processes of infection of *C. fructicola*. KEGG pathways significantly enriched only in the Wuzhishan population included ribosome biogenesis in eukaryotes (ko03008), spliceosome (ko03040), RNA transport (ko03013), RNA polymerase (ko03020), and purine metabolism (ko00230). Pathways that were significantly enriched only in the Shaoyang group included phenylalanine metabolism and beta-alanine metabolism.

**Table 4.** KEGG pathways significantly enriched in oil tea leaves inoculated with strains of *C. fructicola* and related up-regulated genes.

| Pathway ID | Pathway | All genes with pathway annotation | DEGs with pathway annotation |  | Qvalue |  |
| --- | --- | --- | --- | --- | --- | --- |
|  |  |  | WZS | SY | WZS | SY |
| ko03010 | Ribosome | 102 | 99 | 88 | 7.04E-30 | 2.34E-20 |
| ko00330 | Arginine and proline metabolism | 79 | 51 | 50 | 3.45E-03 | 1.07E-03 |
| ko00230 | Purine metabolism | 99 | 62 | 37 | 3.10E-03 | 0.9999 |
| ko03008 | Ribosome biogenesis in eukaryotes | 71 | 56 | 42 | 9.59E-08 | 0.0231 |
| ko03013 | RNA transport | 93 | 63 | 41 | 9.49E-05 | 0.9079 |
| ko03020 | RNA polymerase | 26 | 22 | 12 | 5.79E-04 | 0.9727 |
| ko03040 | Spliceosome | 88 | 68 | 40 | 1.33E-08 | 0.7774 |

Comparison of differential genes associated with the purine metabolism pathway revealed that 35 differential genes were up-regulated in the Wuzhishan group and Shaoyang group, while the remaining 27 up-regulated genes in the Wuzhishan group were not up-regulated in the Shaoyang group. Figure 5 shown the heatmap of the 27 up-regulated genes specific to the WZS group. Of the up-regulated genes specific to the purine metabolism pathway following inoculation with the Wuzhishan population, 12 of the 27 genes are involved in the regulation of purine biosynthesis and catabolic metabolism (Table 5). Related downstream products of this pathway include guanylate synthase, adenylate deaminase, inosine cyclic hydrolase, and adenine phosphoribosyltransferase. The remaining 17 genes of the Wuzhishan population an up-regulated gene list is involved in RNA and DNA synthesis. Purine biosynthesis includes de novo synthesis and remedy pathway (Figure 7).

**Table 5.** 12 Up-regulated genes specific to the WZS group related to purine synthesis and catabolism.

| Gene ID | Gene | Description | Pathway Module |
| --- | --- | --- | --- |
| CGGC5_8860 | APT1 | Adenine<br>phosphoribosyltransferase | -- |
| CGGC5_5780 | NT5E | 5'-nucleotidase | -- |
| CGGC5_5691 | UAZ | Uricase | Purine degradation |
| CGGC5_3653 | PRS5 | Ribose-phosphate<br>pyrophosphokinase | PRPP biosynthesis |
| CGGC5_3645 | ADE6 | Phosphoribosylformylglycinamid<br>ine synthase | IMP biosynthesis |
| CGGC5_2412 | ADE17 | IMP cyclohydrolase | IMP biosynthesis |
| CGGC5_15250 | ADA1 | AMP deaminase | -- |
| CGGC5_13215 | AAH1 | Adenosine deaminase | -- |
| CGGC5_11685 | GUA1 | GMP synthase | Guanine ribonucleotide<br>biosynthesis |
| CGGC5_11516 | SPCC830.11c | Adenylate kinase | Adenine ribonucleotide<br>biosynthesis |
| CGGC5_11342 | ADE3 | Phosphoribosylformylglycinamid<br>ine synthase, partial | IMP biosynthesis |
| CGGC5_11063 | NCU09789 | Adenylosuccinate synthetase | Adenine ribonucleotide<br>biosynthesis |

**Figure 5.**
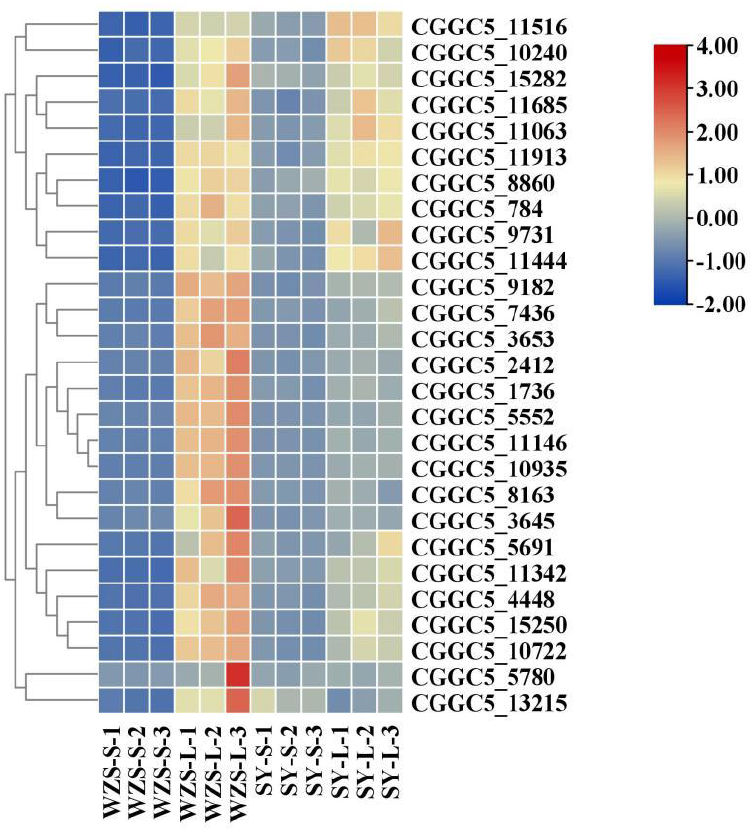
Heatmap of the 27 up-regulated genes specific to the WZS group. The heatmap was created using the TBtool (version 0.66837, (Chen, C., et al, 2018)) function “heatmap” with default parameter setting and shows normalized FPKM values.

**Figure 6.**
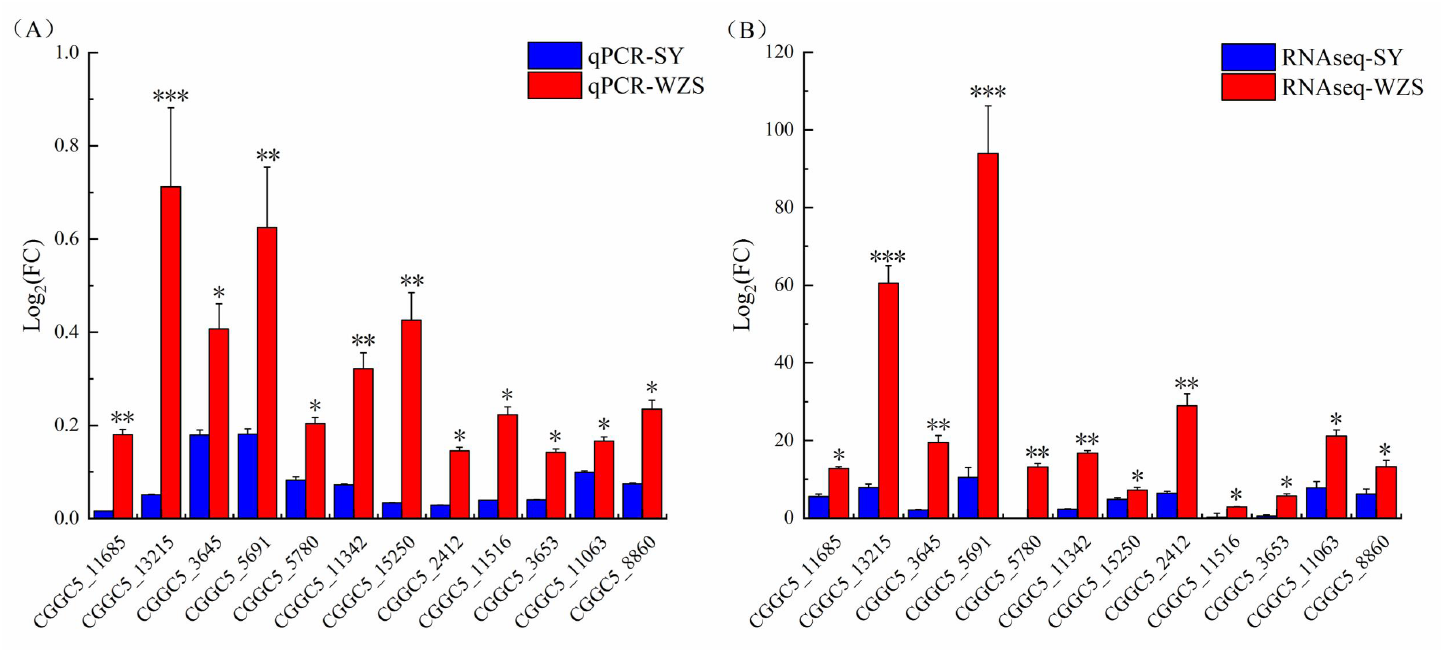
Expression verification of 12 up-regulated genes specific to the WZS group. A: qRT-PCR verification result of WZS-L-vs-SY-L. B: RNAseq verification result of WZS-L-vs-SY-L.

**Figure 7.**
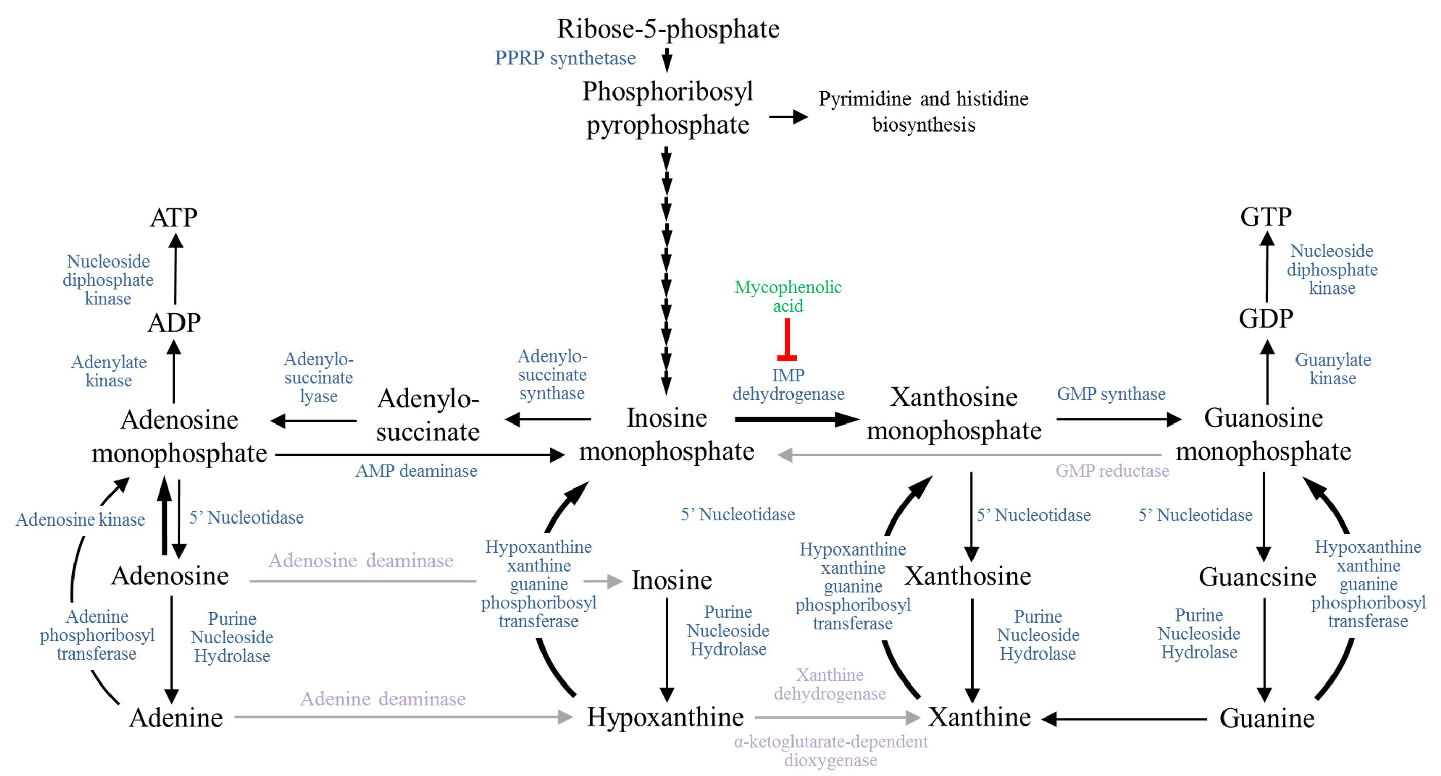
Components of the purine metabolic pathway (Morrow, C. A., et al, 2012). The de novo synthesis pathway uses 5-phosphate ribose (Ribose-5P) as the raw material, and generates phosphoribosyl pyrophosphate (PRPP) via phosphoribosyl pyrophosphate kinase (PRS), which then synthesizes inosinic acid (IMP) through ten consecutive reactions catalyzed by various enzymes. In the de novo synthesis pathway, IMP is converted into adenylate (AMP), or to xanthosine (XMP) and then guanylate (GMP), and finally to ATP and GTP. In the rescue pathway, free adenine and guanine in cells synthesize AMP and GMP under the action of adenine phosphoribosyltransferase (APT). The purine degradation pathway first converts purine nucleotides, such as AMP and GMP, into xanthine by various enzymes. Xanthine is finally degraded to produce uric acid.

## 4 Discussion

### 4.1 *C. fructicola* Collected in Wuzhishan Have a Stronger Ability to Infect Oil Tea Leaf Tissue

The two regions selected for comparison of *C. fructicola* populations are approximately 7 ° in latitude apart. The average annual rainfall and temperature of Wuzhishan is higher than that of Shaoyang. However, Shaoyang experiences greater temperature variation, with winter temperatures which are not suitable for proliferation of *C. fructicola*. Therefore, we speculate that the anthracnose pathogen’s wintering time in Wuzhishan is shorter, and that time of onset is earlier. Therefore, due to the presences of conditions which promote growth of anthracnose, we speculate that annual incidence of anthracnose in *Oil tea* in Wuzhishan is greater and occurs over longer durations of time.

A comparison of biological characteristics and pathogenicity of the two geographical populations of *C. fructicola* found that the number of lesions 96 hours post inoculation(hpi) did not differ statistically. However, there was a significant difference in the diameter of the lesions among the two populations of *C. fructicola* 96 hours after inoculation on detached leaves of oil tea. The spread of lesions on *Oil tea* caused by the Wuzhishan population was more severe, indicating that *C. fructicola* from Wuzhishan had a stronger ability to destroy *Oil tea* leaf tissue.

Based on results of the phenotypic experiments, the two populations of *C. fructicola* had similar abilities to infect the host however their ability to further colonize differed.

### 4.2 Changes in *C. fructicola* Carbon Source Utilization on Oil Tea Leaf Tissues in the Late Infection Stage

RNA sequencing results demonstrated gene, GO term and KEGG pathway enrichment related to ribosomal activity for both populations. Enriched GO cell composition terms included ribosomes in both populations, this result indicates that mitochondria and endoplasmic reticulum ribosomes might play important roles in the progress of infection and illustrates the active regulation of various proteins by pathogenic fungi after infection. Proteins like hexose transporters can be used to absorb nutrients from host cell tissues or to resist defensive immune stress in plants.

After infection by pathogenic fungi, carbohydrates in the apoplast are important carbon sources for pathogenic fungi. Previous work has found that four hexose transporters (CgHxt1, CgHxt2, CgHxt3, and CgHxt5) of *Colletotrichum graminicola* can transport a variety of hexoses, including fructose, mannose, galactose and xylose. The transporters *CgHxt2* and *CgHxt5* are only expressed in the vegetative phase based on dead plant tissue (Lingner, U., et al, 2011). Another study showed that *MFS1* gene knockouts of *Colletotrichum lindemuthianum* had defects in the use of glucose, mannose and fructose. Furthermore, semi-quantitative PCR analysis found that the *MFS1* gene was only up-regulated during the vegetative phase based on dead plant tissue 96 hours after infecting the host (Pereira, M. F., et al, 2013). In the current study, GO functional enrichment analysis suggests that *C. fructicola* transforms and utilizes sugar derived from plants. Galactosidase is a class of enzymes that hydrolyze galactosyl bond-containing substances, and hydrolysis produces substances such as galactose, glucose and fructose (Yan, Q. J., et al, 2017). Although there exist few studies on enzymes related to molecular mechanisms of phytopathogenic fungi, it can be inferred that galactosides were hydrolyzed by *C. fructicola* 96 hours after infection with oil tea as a new carbon source.

### 4.3 Glutamine Family Amino Acids Help *C. fructicola* Resist Host Immune Responses during Late Stages of Infection

Glutamine family amino acids include arginine, proline, glutamic acid, and glutamine, where arginine is a precursor of proline and glutamic acid. A number of studies have demonstrated that synthesis of arginine by fungi has important effects on growth and development, production of conidia, *MoARG1, MoARG5, 6* and *MoARG7* by *Magnaporthe oryzae* (Zhang, Y., et al, 2015), *ARG1* by *Fusarium oxysporum* (Namiki, F., et al, 2001), *PATH-19* and *PATH-35* by *Colletotrichum higginsianum* (Takahara, H., et al, 2001). The anti-stress effects of proline in plants and endophytic fungi has been widely studied (Mona, S. A., 2017, Xu, L., 2017, Liang, X., et al, 2013). A number of plants can accumulate large amounts of proline when exposed to stresses such as reactive oxygen stress, therefore proline is often used as a physiological or biochemical indicators of plant stress resistance (Zhu, H. W., et al, 2009). However, the role of proline in pathogenic fungi has not received much attention. For example, proline has been reported to remove reactive oxygen species in *Colletotrichum truncatum* (Chen, C., Dickman, M. B., 2005).

Glutamine and glutamic acid can be interconverted by enzymes: In *Saccharomyces cerevisiae*, glutamine can be directly converted to glutamate by glutamine enzymes. In addition, glutamine can be degraded or converted to glutamate by NADH-dependent glutamate synthase (Miller, S. M., Magasanik, B., 1990). In addition, glutamine and glutamic acid are involved in biosynthesis of glutathione. Glutathione exists in two forms: an oxidized state (GSSG) or reduced state (GSH). During conversion of GSH to GSSG, its thiol group is used as an electron donor to maintain the activity of thiol proteins and enzymes, and to reduce host-derived ROS stress on pathogen cells. Furthermore, the glutathione antioxidant system is important to pathogens to facilitate infection of hosts, therefore glutamine family amino acids play an important role in resisting plant immune responses.

### 4.4 Up-regulation of Purine Metabolism Improves Growth of *C. fructicola*

By use of GO analysis, it was observed that ribosome-related genes of Wuzhishan and Shaoyang fungi populations were significantly up-regulated following infection. However, the number of differentially expressed genes associated with ribosome-related annotations in the Shaoyang group was significantly less than that in the Wuzhishan group. In addition, comparison of GO analysis results of differentially expressed gene lists demonstrated enrichment in activity of translation factors for the Wuzhishan group but not the Shaoyang group, indicating that the translation factor activity of the Wuzhishan group was significantly higher during the process of ribosomal translation. KEGG enrichment analysis indicated that the Wuzhishan group was significantly enriched in signal pathways such as spliceosome, RNA transport, and RNA polymerase, while they were not significantly enriched in the Shaoyang group. Differences in transcriptional and translational activity of the two populations of *C. fructicola* indicate that the Wuzhishan population are more active, and that enriched pathways and differentially expressed genes might be related to its stronger pathogenicity.

Purine nucleotide metabolism is necessary for biological metabolic processes and expression of genetic information, and supports a number of physiological and biochemical reactions (Zrenner, R., et al, 2006). Purine in living organisms can be divided into adenine, guanine, xanthine and hypoxanthine according to their base pairs. Purines are not only an important part of nucleic acid, but also participate in protein translation, phosphate utilization and energy metabolism (Gauthier, S., et al, 2008). Using KEGG enrichment analysis, we found that differentially expressed genes in the Wuzhishan group were significantly enriched in processes related to purine metabolism, indicating that *C. fructicola* in Wuzhishan group induce purine related genes associated with the nucleotide metabolic pathway after infection. We found that 33 differentially expressed genes in the Wuzhishan group and Shaoyang group were up-regulated following infection, while the remaining 29 of the 62 up-regulated genes were specific to the Wuzhishan group. There were no significant differences in expression levels before and after infection. Twelve of the 29 genes that were specific to the Wuzhishan group were involved in regulation of purine biosynthesis and catabolic metabolism (Table 5), especially biosynthetic pathways. Of the 12 genes, four (*PRS5, ADE3, ADE17, ADE6*) are involved in regulating biosynthesis of IMP in the purine de novo synthesis pathway, three (*ADSS, ADK, GUA1*) are involved in regulating ATP and GTP synthesis by IMP, one (*APT1*) participates in regulation of purine rescue pathways, and four (*AAH1, ADA1, UAZ, NT5E*) participate in regulation of purine degradation pathways.

Genes that were specifically up-regulated in the Wuzhishan group play important roles in purine anabolic metabolism, and data show that the purine de novo synthesis pathway is directly related to the growth and development and pathogenicity of pathogenic fungi (Yemelin, A., et al, 2017). Previously, it has been observed that the guanylate kinase and inosine-5’-phosphate lactate dehydrogenase (IMPDH) play key roles in the GTP synthesis pathway. Knock down of the Guanylate kinase gene, *MoGuk2*, in *Magnaporthe oryzae* led to reductions in the expansion of hyphae by the host (Cai, X., et al, 2017). The five active sites of the IMPDH coding gene, *MoIMD4*, are involved in regulating the pathogenicity of *M. oryzae*, thus *MoIMD4* knockout mutant are less pathogenic than wild-type *M. orzae* (Yang, L., et al, 2019). *ACD1* of *Fusarium graminearum* participates in the regulation of the conversion process of AMP to IMP. The knockout mutant of *ACD1* cannot form ascospores, and the expansion ability of the infection hyphae decreases in the host cells. Further analyses of phenotypes were performed in the knockout mutant of the *APT1, XPT1, AAH1*, and *GUD1* genes, revealed that the growth rate and pathogenicity of these four mutants were not significantly different from those of the wild type, which indicated when a de novo purine synthesis pathway exist in *F. graminearum*, the *APT1, XPT1, AAH1*, and *GUD1* genes involved in regulating the purine salvage pathway are not necessary for the growth and pathogenicity of *F. graminearum* (Sun, M., 2019). Our analysis indicates that representative strains of Wuzhishan *C. fructicola* have increased purine metabolism, which might be related to its greater pathogenicity.

## 7 Acknowledgments

We thank the Guangzhou Genedenovo Biotechnology Company for assisting with the sequencing analysis.

## Reference

Cui, X., Wang, W., Yang, X., Li, S., Qin, S., Rong, J. (2016). Potential distribution of wild Camellia oleifera based on ecological niche modeling. Biodiversity Science, 24(10), 1117–1128.

Zhou, W., Qiang, W., Yang, J., Wang, J., Lian, X., & Xu, L. (2017). Establishment of DNA Fingerprints and Cluster Analysis for Oil Camellia Cultivars Based on SSR Markers. Molecular Plant Breeding, 238–249. (in Chinese)

YE Jian-ren, H.W. (2011). Forest pathology (3rd edition ed.). Beijing: China Forestry Press. (in Chinese)

Chen, X., Wang, J., Sun, S., & Wu, H. X. (2009). Histopathological observation on the invasion of Colletotrichum gloeosporioides to oil tea. Forest Pest Disease.

Wang, H. K., Lin, F. C., & Wang, Z. Y. (2004). Mechanical penetrating force of appressoria of plant pathogenic fungi. mycosystema, 23(1), 151–157.

Jiang, S. Q., & Li, H. (2018). First Report of Leaf Anthracnose Caused by Colletotrichum karstii on Tea-Oil Trees (Camellia oleifera) in China. Plant Disease, 102, PDIS-08-17-1195-PDN.

Li, H., Li, Y., Jiang, S., Liu, J., & Zhou, G. (2017). Pathogen of Oil-Tea Trees Anthracnose Caused by Colletotrichum spp. in Hunan Province. Linye Kexue/Scientia Silvae Sinicae, 53, 43–53.

Tang, Y. L., Zhou, G. Y., & Li, H. (2015). Identification of a new anthracnose of Camellia oleifera based on multiple-gene phylogeny. J. Tropical Crops, 36(5), 972–977.

Li, H. (2018). Population Genetic Analyses of the Fungal Pathogen Colletotrichum on Tea-Oil Trees in China and Characterization of a MAPK gene CfPMK1 in the Pathogen. (Dissertation), Central South University of Forestry & Technology, Changsha, China.

Deng, L., Liu, J. A., Zhou, G. Y., Zhang, N., & He, Y. H. (2017). Antifungal activity of broth of Camellia anthracnose of antagonistic actinomycete CF17 and isolation and purification of its antifungal substances. Fujian Agric. Forest. Univ. (Nat. Sci. Ed.), 46(01), 15–20.

Zhou, Y. H., Zhang, L., Deng, L. L., & Zeng, K. F. (2015). Mechanisms of interaction between Pichia membranaceus-bound chitosan and anthrax-citrus fruits. Paper presented at the the 2015 Annual Conference of the Chinese Society of Plant Pathology, Haikou, China. (in Chinese)

Deng, Y. Z., & Naqvi, N. I. (2019). Metabolic Basis of Pathogenesis and Host Adaptation in Rice Blast. Annual Review of Microbiology, 73, 601–619.

Kenneth JC. (2002). Strawberry Anthracnose: Histopathology of Colletotrichum acutatum and C. fragariae. Phytopathology, 10(92), 1055–1063.

Ma, W. (2010). The evaluation for forests ecosystem purify the atmosphere function in Shaoyang city. J. Central South Univ. Forest. Technol., 30, 51–54.

Su, F. (2018). Study on Germplasm Resources and Antioxidant Activity of Wild Tea Trees in Wuzhishan City. (Master’s thesis), Hainan University, Haikou, China. (in Chinese)

Zheng, D. J., Pan, X. Z., & Zhang, D. M. (2016). Survey and analysis on tea-oil camellia resource in Hainan. J. Northwest Forest. Univ., 31, 130–135.

He, L., Zhou, G. Y., Liu, J. A., & Xu, J. P. (2016). Population Genetic Analyses of the Fungal Pathogen Colletotrichum fructicola on Tea-Oil Trees in China. Plos One, 11(6), e0156841. (in Chinese)

Chen, C., Xia, R., Chen, H., & He, Y. (2018). a Toolkit for Biologists integrating various biological data handling tools with a user-friendly interface. BioRxiv. doi:10.1101/289660

Morrow, C. A., Valkov, E., Stamp, A., Chow, E. W. L., Lee, I. R., Wronski, A., Fraser, J. A. (2012). De novo GTP biosynthesis is critical for virulence of the fungal pathogen Cryptococcus neoformans. PLoS Pathog, 8, e1002957.

Lingner, U., Münch, S., Deising, H. B., & Sauer, N. (2011). Hexose transporters of a hemibiotrophic plant pathogen: functional variations and regulatory differences at different stages of infection. Journal of Biological Chemistry, 286, 20913–20922.

Pereira, M. F., De Araújo Dos Santos, C.M., De Araújo, E. F., De Queiroz, M. V., & Bazzolli, D. M. S. (2013). Beginning to understand the role of sugar carriers in Colletotrichum lindemuthianum: the function of the gene mfs1. J. Microbiol, 51, 70–81.

Yan, Q. J., Liu, Y., & Jiang, Z. Q. (2017). Advances in microbial α-galactosidase. J. Microbiol., 37, 1–9.

Zhang, Y., Shi, H., Liang, S., Ning, G., Xu, N., Lu, J., Lin, F. (2015). MoARG1, MoARG5,6 and MoARG7 involved in arginine biosynthesis are essential for growth, conidiogenesis, sexual reproduction, and pathogenicity in Magnaporthe oryzae. microbiological Research, 180, 11–22.

Namiki, F., Matsunaga, M., Okuda, M., Inoue, I., Nishi, K., & Fujita, Y. (2001). Mutation of an arginine biosynthesis gene causes reduced pathogenicity in Fusarium oxysporum f. sp melonis. Molecular Plant-Microbe Interactions, 14(4), 580–584.

Takahara, H., Huser, A., & O’connell, R. (2012). Two arginine biosynthesis genes are essential for pathogenicity of Colletotrichum higginsianum on Arabidopsis. Mycology, 3, 54–64.

Mona, S. A., Hashem, A., Abd_Allah, E. F., Alqarawi, A. A., Soliman, D. W. K., Wirth, S., & Egamberdieva., D. (2017). Increased resistance of drought by Trichoderma harzianum fungal treatment correlates with increased secondary metabolites and proline content. Journal of Integrative Agriculture, 16, 1751–1757.

Xu, L., Wang, A. A., Wang, J., Wei, Q., & Zhang, W. Y. (2017). Piriformospora indica confers drought tolerance on Zea mays L. through enhancement of antioxidant activity and expression of drought-related genes. The Crop Journal, 5, 251–258.

Liang, X., Zhang, L., Natarajan, S. K., & Becker., D.F. (2013). Proline Mechanisms of Stress Survival.. Antioxidants & Redox Signaling., 19, 998–1011.

Zhu, H.W.W.; Yan, Y. (2009). Effect of proline on plant growth under different stress conditions. J. Northeast Forest. Univ., 37, 86–89.

Chen, C., & Dickman, M. B. (2005). Proline suppresses apoptosis in the fungal pathogen Colletotrichum trifolii. Proceedings of The National Academy of Sciences of The U.S.A., 102, 3459–3464.

Miller, S. M., & Magasanik, B. (1990). Role of NAD-linked glutamate dehydrogenase in nitrogen metabolism in Saccharomyces cerevisiae. J. Bacteriol., 172, 4927–4935.

Zrenner, R., Stitt, M., Sonnewald, U., & Boldt, R. (2006). Pyrimidine and purine biosynthesis and degradation in plants. Annu. Rev. Plant Biol., 57, 805–836.

Gauthier, S., Coulpier, F., Jourdren, L., Merle, M., Beck, S., Konrad, M., Pinson, B. (2008). Co-regulation of yeast purine and phosphate pathways in response to adenylic nucleotide variations. Mol. Microbiol, 68, 1583–1594.

Yemelin, A., Brauchler, A., Jacob, S., Laufer, J., Heck, L., Foster, A., Andresen, K. T. E. (2017). Identification of factors involved in dimorphism and pathogenicity of Zymoseptoria tritici.. Plos One, 12(8), e0183065.

Cai, X., Zhang, X., Li, X., Liu, M., Liu, X., Wang, X.,Zhang, Z. (2017). The atypical guanylate kinase MoGuk2 plays important roles in asexual/sexual development, conidial septation, and pathogenicity in the rice blast fungus. Front. Microbiol, 8, 2467.

Yang, L., Ru, Y., Cai, X., Yin, Z., Liu, X., Xiao, Y., Zhang, Z. (2019). MoImd4 mediates crosstalk between MoPdeH-cAMP signalling and purine metabolism to govern growth and pathogenicity in Magnaporthe oryzae. Mol. Plant Pathol, 20, 500–518.

Sun, M. (2019). Characterization of mRNA Splicing Related SNU66 and Purine Nucleotide Metabolism Pathway Genes in Fusarium Graminearum. (Dissertation), Northwest A & F University, Yangling, Shaanxi, China. (in Chinese)

